# Cryo-EM single particle structure refinement and map calculation using *Servalcat*

**DOI:** 10.1101/2021.05.04.442493

**Authors:** Keitaro Yamashita, Colin M. Palmer, Tom Burnley, Garib N. Murshudov

## Abstract

In 2020, cryo-EM single particle analysis achieved true atomic resolution, thanks to technological developments in hardware and software. The number of high resolution reconstructions continues to grow, increasing the importance of accurate determination of atomic coordinates. Here, a new Python package and program called *Servalcat* is presented that is designed to facilitate atomic model refinement. *Servalcat* implements a refinement pipeline, using the program *REFMAC*5 from the *CCP4* package. After the refinement, *Servalcat* calculates a weighted *F*_o_ − *F*_c_ difference map, which was derived from Bayesian statistics. This map helps manual and automatic model building in real space, as is common practice in crystallography. The *F*_o_ − *F*_c_ map helps visualisation of weak features including hydrogen densities. Although hydrogen densities are weak, they are stronger than in electron density maps produced by X-ray crystallography, and some hydrogen atoms are even visible at ∼ 1.8 Å resolution. *Servalcat* also facilitates atomic model refinement under symmetry constraints. If a point group symmetry has been applied to the map during reconstruction, the asymmetric unit model is refined with appropriate symmetry constraints.

## 2. Introduction

Atomic model refinement is the optimisation of the model’s parameters against observed data. Atomic parameters typically include coordinates, atomic displacement parameters (ADPs), and occupancies. In crystallography, refinement is crucial because of the phase problem: the accuracy of density maps relies on the accuracy of the phases of the structure factors. Accurate phases are not observed and must be calculated from the model (Tronrud, 2004). More accurate maps may be obtained as the model becomes more accurate through the refinement. In single particle analysis (SPA) there is no phase problem, although the Fourier coefficients can be noisy, especially at high resolution.

Accurate atomic model determination is becoming more and more important due to the “resolution revolution” in cryo-EM SPA following the introduction of direct electron detectors and new data processing methods (Bai *et al*., 2015). As of April 2021, there are more than 2,500 SPA entries with resolution better than 3.5 Å deposited in the Electron Microscopy Data Bank (EMDB) (Tagari *et al*., 2002). This improvement in resolution has accelerated the development of methods for model building, refinement and validation. Automatic model building programs developed originally for crystallography are now being adapted for cryo-EM SPA maps (Terwilliger *et al*., 2018*a*; Hoh *et al*., 2020; Chojnowski *et al*., 2021). Density modification and local map sharpening can help interpret the map (Jakobi *et al*., 2017; Terwilliger *et al*., 2018*b*; Ramírez-Aportela *et al*., 2019; Ramlaul *et al*., 2019; Terwilliger *et al*., 2020). In general, care must be exercised when using any techniques based on prior knowledge; bias towards incorrect assumptions might lead to misinterpretation of the maps. Full-atom refinement can be done either in real space (Afonine *et al*., 2018) or in reciprocal space (Murshudov, 2016).

After refinement, the model should be validated; the model should have a reasonable geometry and should describe the map well. Due to the low data-to-parameter ratio, all models will exhibit a degree of overfitting; however the model should not deviate substantially from cross-validation data (Brown *et al*., 2015). *MolProbity* is the most widely used geometry validation tool, which includes analyses of clashes, rotamers, and the Ramachandran plot (Chen *et al*., 2010). Map-model quality is assessed using real space local correlations (Cragnolini *et al*., 2021), which have been commonly used in crystallography (Tickle, 2012). In reciprocal space refinement, the *R* factor can be calculated as in crystallography, but the map-model Fourier shell correlation (FSC) is preferred as it does not depend on resolution-dependent scaling and takes phases into account explicitly. An *F*_o_ − *F*_c_ map, which highlights unmodelled features and errors in the current model, is almost always used in crystallography, and some similar tools already exist for SPA (Joseph *et al*., 2020). The *σ*_*A*_-weighted 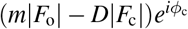 map as used in crystallography is not directly applicable to SPA, because phases are available for both *F*_o_ and *F*_c_ and we should model the error of *F*_o_ in the complex plane, rather than simply using the estimated phase error as in crystallography (see below).

In 2020, cryo-EM SPA achieved atomic resolution – according to Sheldrick’s criterion (Wlodawer & Dauter, 2017) – in structural analyses of apoferritin, which were reported by two groups (Nakane *et al*., 2020; Yip *et al*., 2020). Nakane *et al*. (2020) observed hydrogen atom densities at 1.2 and 1.7 Å resolutions using *F*_o_ − *F*_c_ maps calculated by *REFMAC*5. There is a higher chance of observing hydrogen density in electron microscopy than in X-ray crystallography, because of the increased contrast for the lighter elements (Clabbers & Abrahams, 2018). Nevertheless, hydrogen density is relatively weak and there is always a much higher peak from the parent atom nearby, so the *F*_o_ − *F*_c_ difference map is essential to see them. In addition, there is complexity in the interpretation of hydrogen peaks in EM. An electron in a hydrogen atom is usually shifted towards the parent atom from the nucleus position. In EM, both the electrons and the nucleus contribute to scattering, and this offset results in a shift of hydrogen density peaks beyond the position of the hydrogen nucleus (Nakane *et al*., 2020).

SPA structures often have point group symmetries (rather than space group symmetry as in crystallography). Approximately half of the SPA entries in the EMDB have non-*C*1 point group symmetry according to their associated metadata. Such symmetry is advantageous and helps to reach higher resolution because it increases the effective number of particles. If the map is symmetrised, downstream analyses should be aware of it and the structural model must follow the symmetry. As in crystallography, it is natural to work in a single asymmetric unit. The `MTRIX` records in the PDB format or `struct ncs oper` in the mmCIF format can be used to encode the sy mmetry information^**1**^. Currently, for structures from SPA there are only a few depositions of such asymmetric unit models to the PDB (excepting viruses). We recommend refining and depositing an asymmetric unit model, which makes sure the symmetry copies are truly identical. It should be noted that validation tools must be aware of any applied symmetry operators, but results should be reported for the asymmetric unit only. These considerations are only valid if the map is symmetrised, and we suggest that the point group information should be required by the deposition system.

Here we present *Servalcat*, a Python package and standalone program for the refinement and map calculation of cryo-EM SPA structures. *Servalcat* takes unsharpened and unweighted half maps of the independent reconstructions as inputs, and implements a refinement pipeline using *REFMAC*5, which uses a dedicated likelihood function for SPA (Murshudov, 2016). After the refinement, *Servalcat* calculates a sharpened and weighted *F*_o_ − *F*_c_ map, derived from Bayesian statistics as described below. If the map has a point group symmetry, the user can give an asymmetric unit model and a point group symbol, and the program will output a refined asymmetric unit model with symmetry annotation as well as a symmetry-expanded model. The non-crystallographic symmetry (NCS) constraint function in *REFMAC*5 has been updated to consider symmetry-related non-bonded interactions and ADP similarity restraints (to ensure similarity of ADPs of atoms brought into close proximity via symmetry operations).

*Servalcat* is freely available as a standalone package and also as part of *CCP-EM* (Burnley *et al*., 2017), where the *REFMAC*5 interface has been updated to use *Servalcat*.

## 3. Map calculation and sharpening using signal variance

Let us assume that *F*_o_ is the result of a position-independent blurring *k* of the true Fourier coefficients *F*_T_ with an independent zero-mean Gaussian noise with variance 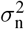. That is,

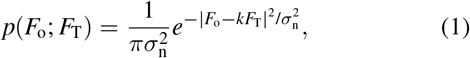

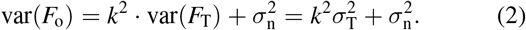

Note that in this work, we treat *k* as a function of resolution *s* . Multiplication by *k* in Fourier space is equivalent to isotropic blurring by a convolution in real space. In general, *k* could take on a different value at each point *s* in Fourier space, which would produce a position-independent but direction-dependent blurring in real space.

The variance of the noise 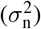 can be calculated from the half maps in resolution bins (Murshudov, 2016).

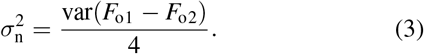

We will later use the relationship of 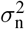 and 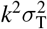 to the FSC, correlation coefficients in resolution bins (Rosenthal & Henderson, 2003):

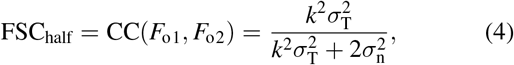

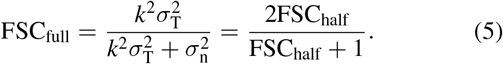

Let us also assume that the errors in the model follow a Gaussian distribution (Luzzati, 1952),

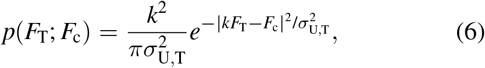

We need two functions; the likelihood *p*(*F*_o_; *F*_c_) for estimation of parameters (of the atomic model and of the distribution function), and the posterior distribution *p*(*F*_T_; *F*_o_, *F*_c_) of the unknown *F*_T_ for map calculation.

### 3.1. Likelihood

As derived in Murshudov (2016),

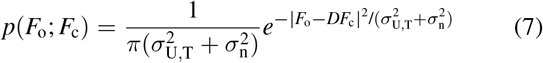

is the likelihood function that is optimized during atomic model refinement. *D* and 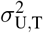 are obtained in each resolution bin *i* by maximising the joint likelihood (7):

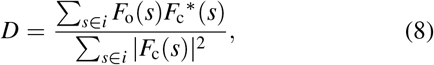

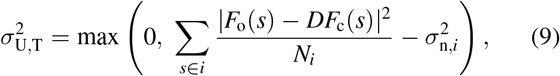

where *N*_*i*_ is the number of Fourier coefficients in bin *i*.

### 3.2. Posterior distribution and map calculation

The posterior distribution, as derived in Murshudov (2016),

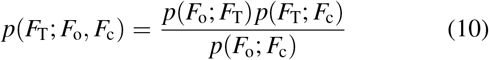

is a 2D Gaussian distribution with the mean and variance

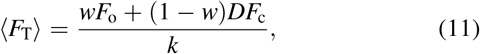

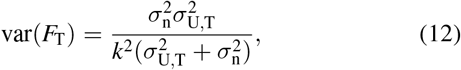

Where

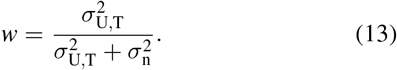

Coefficients for an *F*_o_ − *F*_c_ type difference map can be derived as

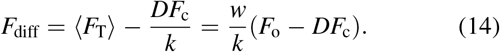

The remaining unknown variable is *k*, which cannot be determined from data alone. For position-independent isotropic Gaussian blurring, *k* has a form of 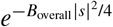, and *B*_overall_ may be estimated from line fitting of a Wilson plot (Wilson, 1942).

However such an estimate is unstable, especially when only low resolution data are available. Here we introduce a simple approximation using the variance of the signal. Let us assume the true map consists of atoms with the same isotropic ADP of ⟨*B*⟩, and then,

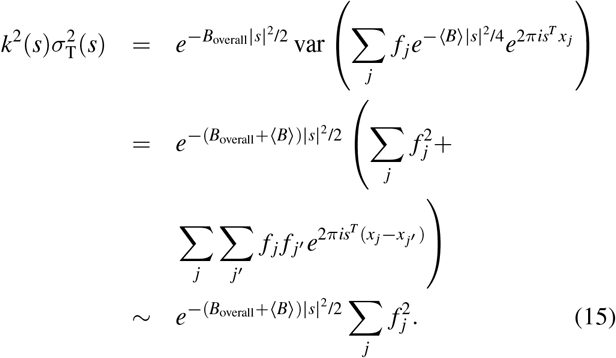

We ignored interference terms 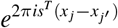. Further ignoring resolution dependent terms in 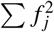, we can use *kσ*_T_ as a proxy for *k*, which gives the best sharpening for the region having an ADP value of ⟨*B*⟩. *kσ*_T_ can be transformed as follows.

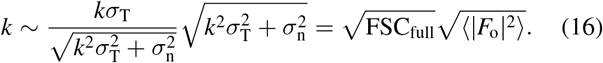

Then the *F*_o_ − *F*_c_ coefficient finally has the form

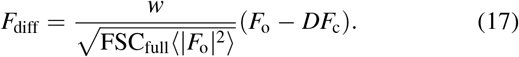

*Servalcat* calculates an *F*_o_ − *F*_c_ map using (17). The significance of difference map peaks is usually defined by r.m.s.d. (sigma) level in crystallography. However, in SPA the box size is arbitrary and the voxels outside the molecular envelope lead to underestimation of the r.m.s.d. value (see the Appendix A). When a mask file is given, *Servalcat* writes a map file after normalizing values within the mask. (Otherwise only the *F*_o_ − *F*_c_ structure factors are written, in MTZ format.) Note that the *F*_o_ − *F*_c_ map is only sensible when the ADPs are properly refined; otherwise we will see peaks due to incorrect ADPs. For that reason, unsharpened *F*_o_ should be used as the input for atomic model refinement (see 4.1); then sharpening is always consistent as the same sharpening factor is applied to *F*_o_ and *F*_c_.

The map from the estimated true Fourier coefficients (11) may be useful, but there is a risk of model bias because of the contribution from *F*_c_. In future, techniques may be available to resolve the issue of model bias. At the moment, *Servalcat* provides the following as a default map for manual inspection. This is a special case of (11) in the absence of model, that is, with *D* = 0,

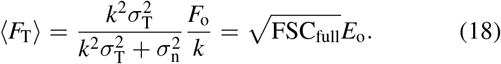

This is equivalent to *EMDA*’s “best map” (https://emda.readthedocs.io).

The approach here should work at any resolution where atomic model refinement is applicable.

## 4. Refinement procedure

In this section the refinement and map calculation procedures are described. Everything other than *REFMAC*5 itself is implemented in *Servalcat* using the *GEMMI* library (https://github.com/project-gemmi/gemmi). Fig. 1 summarizes the procedure.

**Figure 1.**
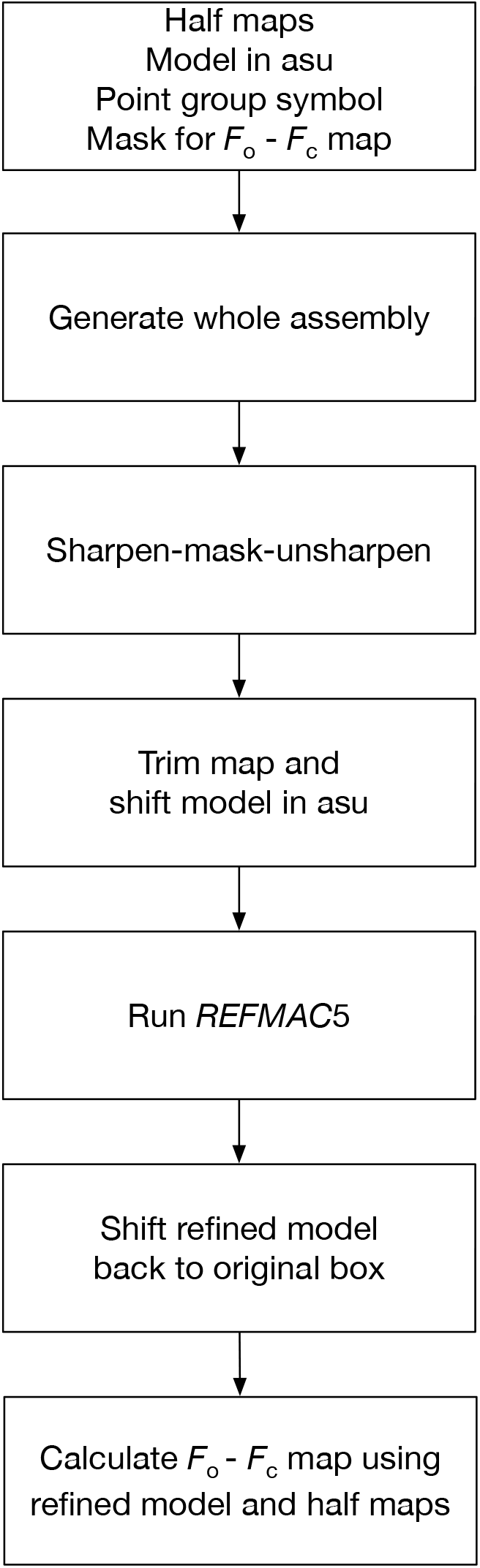
The workflow of *Servalcat* for the refinement of SPA structures.

### 4.1. Map choice

The optimal map depends on the purpose. For manual inspection, optimally sharpened and weighted maps should be used so that the best visual interpretability is achieved. In general, this does not mean the best signal to noise ratio, but it does mean that the details of structural features are visible in the map. On the other hand, unsharpened and unweighted maps are preferred in the refinement. If a sharpened map is used, some atoms may need to be refined to have negative ADPs, but the ADPs are constrained to be positive in the refinement resulting in suboptimal atomic models. On the other hand, blurred maps will just give a shifted distribution of refined ADP values. An unweighted map is preferred because it enables calculation of many properties including noise variance and optimally-weighted maps after refinement (see section 3). Users should therefore be aware that the ADPs in the model are not refined against the same map that is used for visual inspection. Cross-validation (Brown *et al*., 2015) can also be carried out throughout refinement and model building if both half maps are readily available. Therefore, unsharpened and unweighted half maps from two independent reconstructions are considered optimal inputs for the *Servalcat* pipeline, which performs atomic model refinement followed by map calculation.

### 4.2. Masking and trimming

The box size in SPA is often substantially larger than molecule, which is unnecessary for atomic model refinement. Therefore the map is masked and trimmed into a smaller box to speed up calculations as discussed in Nicholls *et al*. (2018). Half maps are first sharpened, masked at a radius of 3 Å (default) from the atom positions, and then blurred by the same factor. Sharpening before masking is important to avoid masking away any of the signal (the tails of the atomic density distributions), because the raw half maps are blurred and the signal is spread out. The optimal sharpening would differ depending on the region, but here we use an overall *B* value estimated by comparing |*F*_o_| with |*F*_c_| calculated from a copy of the initial model with all ADPs set to zero. Alternatively a user-supplied *B* value can be used. The sharpened-masked-unsharpened half maps are then averaged to make a full map that is used as the refinement target in *REFMAC*5. After refinement, the map-model FSC is calculated using a newly created mask based on the refined model.

### 4.3. Point group symmetry

If the maps were symmetrised, the user can specify a point group symbol and give the coordinates for just a single asymmetric unit. Symmetry operators are calculated from the symbols (*Cn, Dn, O, T* , and *I*) following the common orientation convention (Heymann *et al*., 2005). It is also assumed that the centre of the box is the origin of symmetry. This requires translation for each rotation *R*_*j*_, which can be calculated as *c − R*_*j*_*c* = *c*(*I − R*_*j*_) where *c* is the origin of symmetry. Reconstruction programs such as *RELION* (Scheres, 2012) usually follow this assumption. However, the rotation of the axes and position of the origin are arbitrary in general, and in future will be determined automatically using *ProSHADE* (Nicholls *et al*., 2018; Tykac, 2018) and *EMDA*. The model in the asymmetric unit is expanded when creating a mask and performing the map trimming. The rotation matrices are invariant to changing the box sizes and shifts of the molecule. The translation vectors in the symmetry operators are recalculated for the shifted model.

*REFMAC*5 internally generates symmetry copies when calculating *F*_c_ and restraint terms. For anisotropic ADPs, the *B*_aniso_ matrix in the Cartesian basis is transformed by 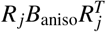. This anisotropic ADP transformation is also implemented in *GEMMI*.

During the refinement, non-bonded interaction and ADP similarity restraints are evaluated using the symmetry-expanded model, and the gradients are calculated for the model in the asymmetric unit.

If atoms are on special positions (e.g. on a rotation axis), they are restrained^**2**^ to sit on the special position and have anisotropic ADPs consistent with symmetry. First, atoms are identified as being on a special position if the following condition is obeyed for any of the symmetry operators *j*:

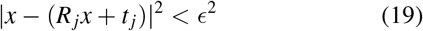

where *ϵ* is a tolerance that can be modified by users. The default value is 0.25 Å . If an atom is on a special position then the program makes sure that symmetry operators for this position form a group that is a subgroup of the point group of the map. Once the elements of the subgroup for this atom have been identified, the atom is forced to be on that position by simply replacing its coordinates with:

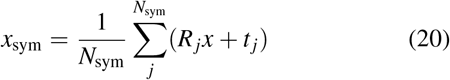

At every cycle, the positions of these atoms are restrained to be on their special positions by adding a term to the target function:

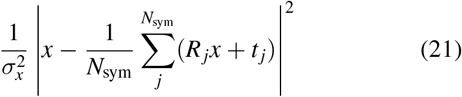

where the summation is performed over all subgroup elements of the special position and *σ*_*x*_ is a user controllable weight parameter for special positions. The occupancy of the atom is adjusted based on the multiplicity of the position.

Anisotropic ADPs of atoms on the special positions are also forced to obey symmetry conditions by replacing the anisotropic tensor with:

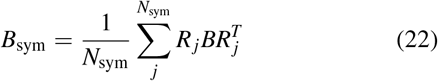

After this, similarly to the positional parameters, at every cycle restraints are applied to the anisotropic tensor of the atoms on special positions to avoid violation of the symmetry condition for the ADP:

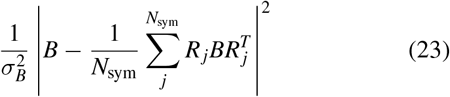

where *σ*_*B*_ is a user controllable weight parameter for *B* values on special positions. Here, the distance between anisotropic tensors is a Frobenius distance |*B*_1_ − *B*_2_|^2^ = Σ_*i, j*_ |*B*_1,*i, j*_ − *B*_2,*i, j*_|^2^.

### 4.4. Hydrogen atoms

Hydrogen electrons are usually shifted towards the parent atoms by 0.1–0.2 Å (Williams *et al*., 2018). This must be accounted for when calculating structure factors from the atomic model (*F*_c_). *REFMAC*5 and *Servalcat* (*GEMMI*) use the Mott-Bethe formula (Mott & Bragg, 1930; Bethe, 1930; Murshudov, 2016) which can conveniently take this fact into account.

The atomic scattering factor for an atom with a shifted nucleus is

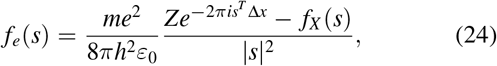

where Δ*x* is the positional shift of the nucleus with respect to the centre of the electron density. The hydrogen density peak in real space is shifted beyond the position of the hydrogen nucleus and varies depending on the ADP and resolution cutoff (Nakane *et al*., 2020). The expected peak position may be calculated by the Fourier transform of (24). The new *CCP4* monomer library includes nucleus bond distances (chem comp bond.value dist nucleus).

### 4.5. Refinement

*REFMAC*5 performs a maximum-likelihood refinement against the Fourier transform of a sharpened-masked-unsharpened map (see 4.2) using a dedicated likelihood function for SPA (7). The estimated noise 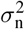 is not used at the moment. No solvent model is used. The average of map-model FSC weighted by the number of Fourier coefficients in each shell (FSC average) is reported to monitor the refinement. At low resolution the use of jelly body restraints or external restraints is encouraged to ensure a large radius of convergence and stabilise the refinement (Murshudov *et al*., 2011; Nicholls *et al*., 2012). Note that jelly body restraints are only useful when the initial model geometry is of good quality because they try to keep the model in its current conformation. After the refinement, *Servalcat* shifts the model back to the original box and adjusts the translation vectors of the symmetry operators if needed. It also generates an MTZ file of map coefficients including the sharpened and weighted *F*_o_ − *F*_c_ and *F*_o_ maps (as calculated by equations (17) and (18)).

### 4.6. User interface

*Servalcat* has a command-line interface. A graphical interface will be available in *CCP-EM*, where the *REFMAC*5 interface has been updated and is now based on *Servalcat*.

From the user’s point of view, the main difference in setting up a refinement job is that the default input is now a pair of half maps. (Refinement from a single input map is still possible but is no longer the default option.) The user is also offered more control over options for refinement weight, symmetry and handling of hydrogen atoms. At the end of refinement, the *F*_o_ − *F*_c_ difference map from *Servalcat* is made available along with the other output files in the *CCP-EM* launcher.

## 5. Methods and results

### 5.1. *F*_o_ − *F*_c_ map for ligand visualisation

*F*_o_ − *F*_c_ omit maps are widely used to convincingly demonstrate the existence of ligands in crystallography. They are also useful for this purpose in SPA. Fig. 2 shows an example of an *F*_o_ − *F*_c_ omit map for the ligand density from EMD-22898 (Kern *et al*., 2021) and EMD-8123 (Murray *et al*., 2016), clearly showing support for the presence of the ligand. To generate the map from EMD-22898, chain A of the atomic model from PDB entry 7kjr was refined using the half maps under *C*2 symmetry constraints. For EMD-8123, PDB entry 5it7 was refined using the half maps without symmetry constraints. After the refinement, the ligand and water atoms were omitted and the *F*_o_ − *F*_c_ maps were calculated. Map values were normalized within a mask. Since a suitable mask for EMD-22898 was not available in the EMDB, one was calculated from half map correlation using *EMDA*.

**Figure 2.**
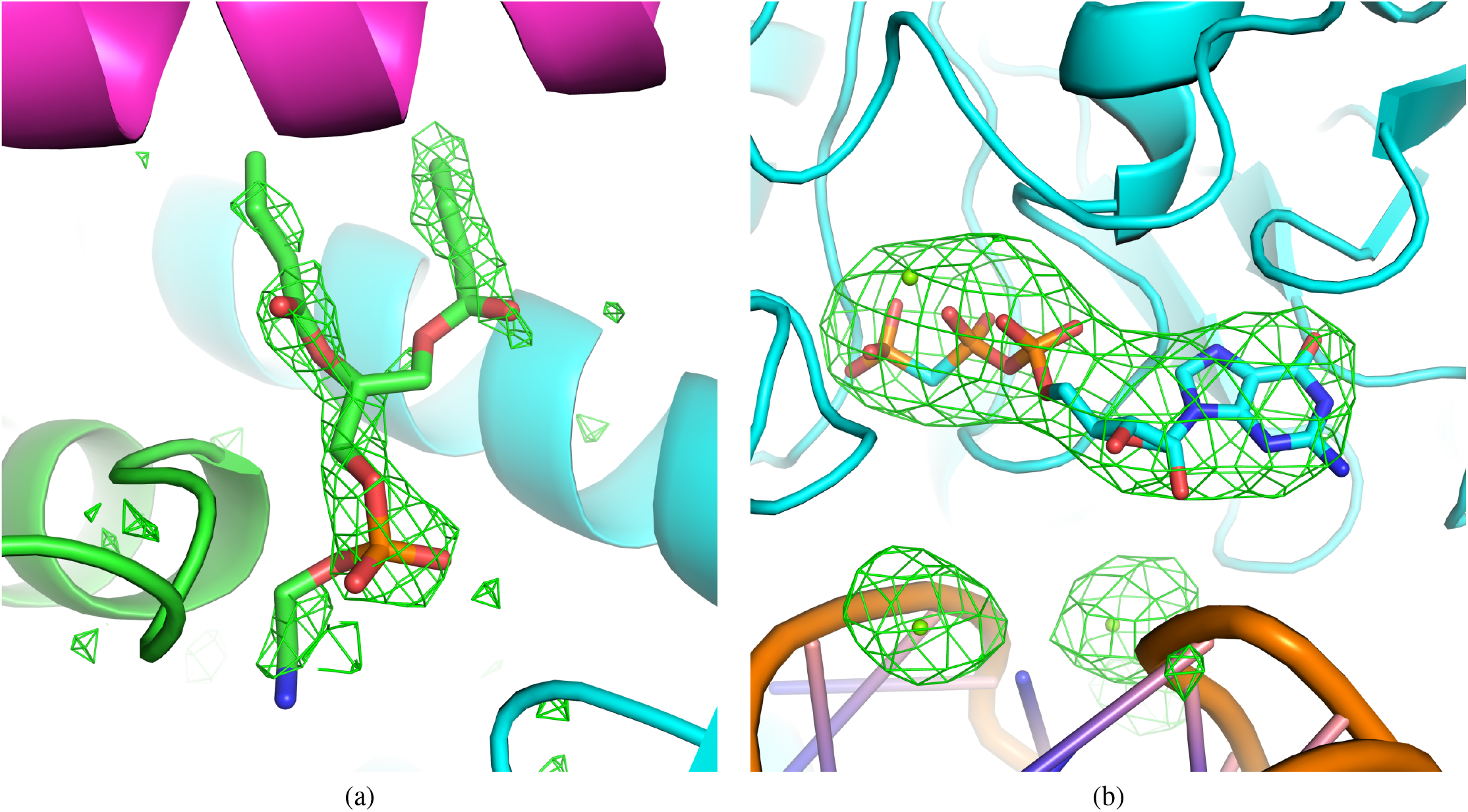
An example of an *F*_o_ − *F*_c_ omit map for visualization of ligand density. The ligand molecules and ions shown as sticks and spheres, respectively, are omitted in the map calculation. The resolution is (a) 2.08 Å (PDB 7kjr/EMD-22898) and (b) 3.6 Å (PDB 5it7/EMD-8123). The *F*_o_ − *F*_c_ omit maps are contoured at 3*σ* (where *σ* is the standard deviation within the mask; see Appendix). The images were created using PyMOL (Schrödinger, LLC, 2020).

*Servalcat*’s weighting and sharpening scheme was compared with alternatives using no weights or 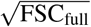 weights (Rosenthal & Henderson, 2003), both with sharpening by the overall *B* value as determined from Wilson plot fitting by *RELION* (Figs. S1 and S2). Especially in the case of EMD-8123 (Fig. S2), sharpening by the overall *B* value obtained by line fitting gave oversharpened maps.

### 5.2. *F*_o_ − *F*_c_ map for detecting model errors

In crystallography, *F*_o_ − *F*_c_ maps are almost always used for manual and automatic model rebuilding. Strong negative density usually indicates that parts of the model should be moved away or removed while strong positive density implies there are unmodelled atoms. The *F*_o_ − *F*_c_ map is typically updated after every refinement session, and refinement may be stopped when there are no significant strong peaks.

The same refinement practice is possible in SPA. Fig. 3 illustrates the use of the *F*_o_ − *F*_c_ map for detecting model errors using EMD-0919 and PDB 6lmt (Demura *et al*., 2020). Chain A of the model was refined using the half maps under *C*8 symmetry constraints. After refinement, the *F*_o_ − *F*_c_ map was calculated and normalized using the standard deviation of the region within the EMDB-deposited mask. In this example it is clear from the positive and negative difference peaks that the tryptophan and methionine side chains should be repositioned. The weighting and sharpening scheme are compared in Fig. S3, demonstrating that appropriate weighting can increase the interpretability of maps.

**Figure 3.**
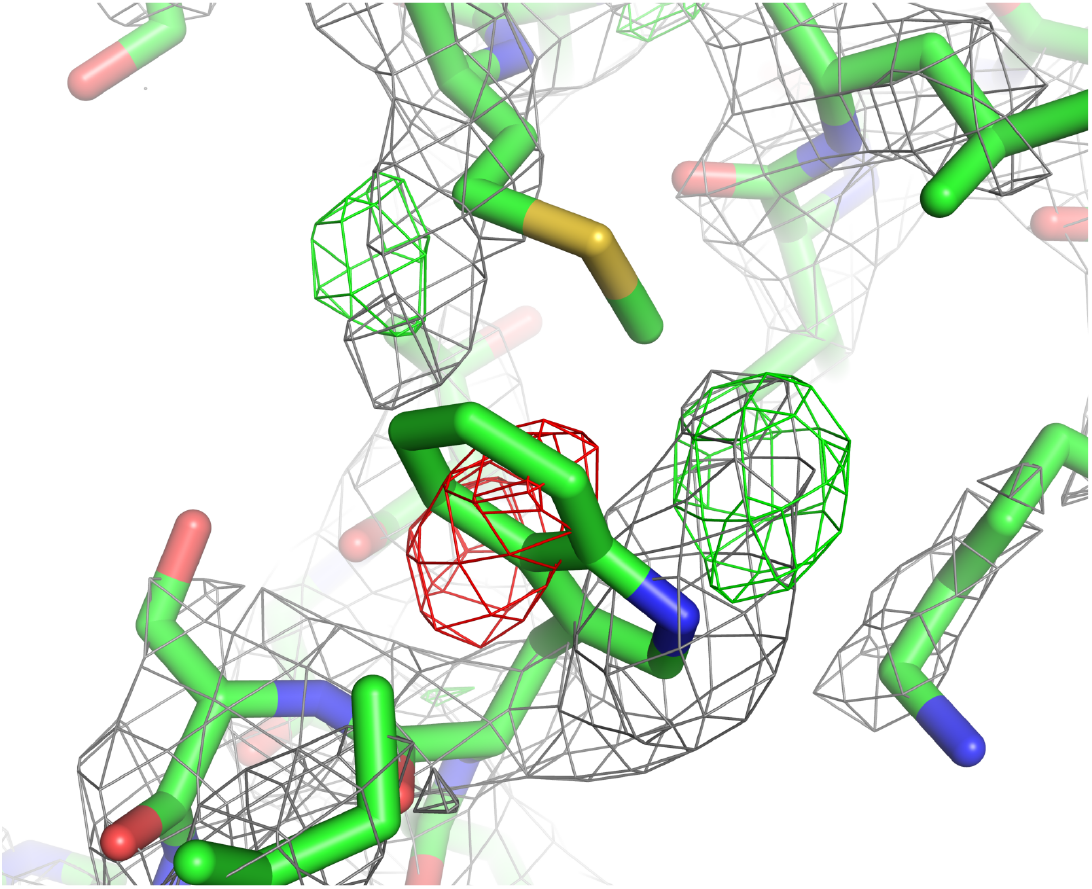
An example of an *F*_o_ − *F*_c_ map for detecting model error, in this case mispositioned tryptophan and methionine side chains (PDB 6lmt/EMD-0919). The resolution is 2.66 Å and the *F*_o_ − *F*_c_ map is contoured at *±*4*σ* (scaled within the mask). Green and red meshes represent positive and negative maps, respectively. Grey mesh is the weighted and sharpened *F*_o_ map. The image was created using PyMOL.

### 5.3. Hydrogen density analysis

Nakane *et al*. (2020) reported convincing densities for hydrogen atoms in apoferritin and GABA_A_R maps by cryo-EM SPA at 1.2 and 1.7 Å resolutions respectively. It is natural to ask what is the lowest resolution at which hydrogen atoms can be seen in cryo-EM SPA using currently available computational tools.

Here, we analyzed apoferritin maps from the EMDB to see if and when hydrogen densities could be observed. There are 25 mouse or human apoferritin entries at resolutions better than 2.1 Å , of which 19 had half maps and were used in the analysis (Table 1). Chain A of each model was refined using the half maps under *O* symmetry constraints. If there was no corresponding PDB code, 7a4m or 6z6u were placed in the map using *MOLREP* (Vagin & Teplyakov, 1997) followed by jigglefit in *Coot* (Brown *et al*., 2015) before full atomic refinement. After 10 cycles of refinement with *REFMAC*5, an *F*_o_ − *F*_c_ map was calculated and normalized within the mask. Riding hydrogen atoms were used in the refinement (so they are not refined, but generated at fixed positions; this is the default in *REFMAC*5) and they were omitted for *F*_o_ − *F*_c_ map calculation. Peaks ≥ 2*σ* and ≥ 3*σ* were detected using *PEAKMAX* from the *CCP4* package (Winn *et al*., 2011), and associated with hydrogen positions if the distance from the peak was less than 0.3 Å . Hydrogen atoms having multiple potential minima (like those in hydroxyl, sulfhydryl or carboxyl groups) were ignored in the analysis. The ratios of the number of hydrogen peaks to the number of hydrogen atoms in the model are plotted in Fig. 4a. The result shows that 1.25 Å data gave the highest ratio of ∼ 70% hydrogens detected (Fig. 5a). Even at 1.84 Å approximately 17% of the hydrogen atoms may be found (Fig. 5b) while at 2.0 or 2.1 Å only a few hydrogen atoms are visible in the map (Fig. 5c). The weighting and sharpening scheme are compared in Figs. S4–6. Note that there may be false positives due to, for example, alternative conformations or inaccuracies in the model.

**Table 1.**
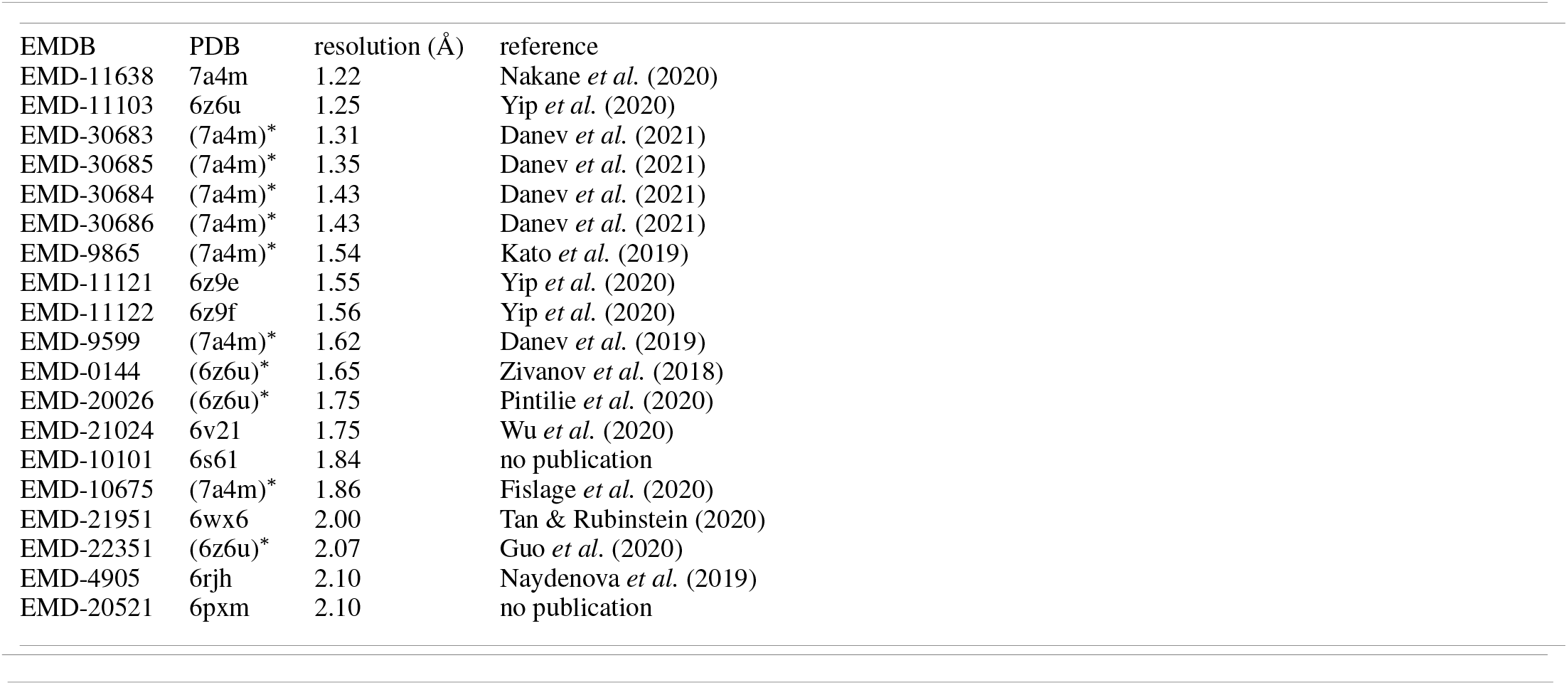
Test data for hydrogen peak analysis. ^∗^No PDB entry was assigned and the code in parenthesis was used for refinement (7a4m from mouse and 6z6u from human).

**Figure 4.**
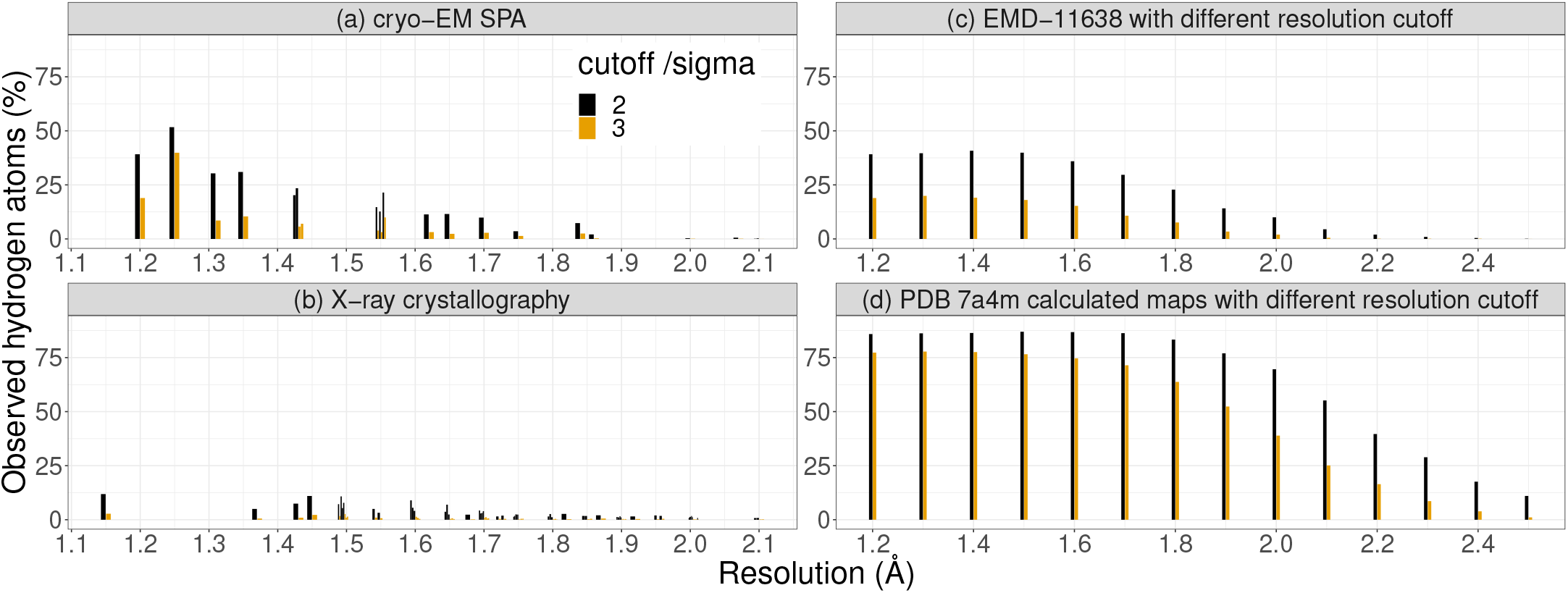
Detection of hydrogen atoms, measured by the number of observed hydrogen density peaks divided by the number of hydrogen atoms in the model. (a) different apoferritin cases by cryo-EM SPA (see Table 1). (b) different (apo)ferritin cases by X-ray crystallography using PDB codes 2v2p, 2v2s, 6gxj, 5erj, 5mij, 2cih, 2w0o, 7bd7, 3f37, 2v2n, 1h96, 2chi, 2zg8, 2v2m, 2z5p, 3h7g, 3f34, 2zg7, 3f32, 3f33, 3f36, 2gyd, 3o7s, 1xz1, 1xz3, 2cn7, 2zg9, 3f38, 2cei, 2iu2, 3fi6, 6env, 3f39, 5ix6, 2v2o, 2v2l, 2v2r, 3o7r, 3rav, 3u90, 3f35, 1aew, 5mik, 2g4h, 2v2i, 3rd0, 5erk, 6ra8, 1gwg, 2clu, 2z5q. (c,d) apoferritin cases calculated at different resolutions from the same map and model, 7a4m/EMD-11638 determined at 1.22 Å . (c) shows detection of hydrogen atoms in *F*_o_ − *F*_c_ maps and (d) in calculated *F*_c_ maps. This figure was prepared using the *ggplot2* (Wickham, 2016) in *R* (R Core Team, 2020).

**Figure 5.**
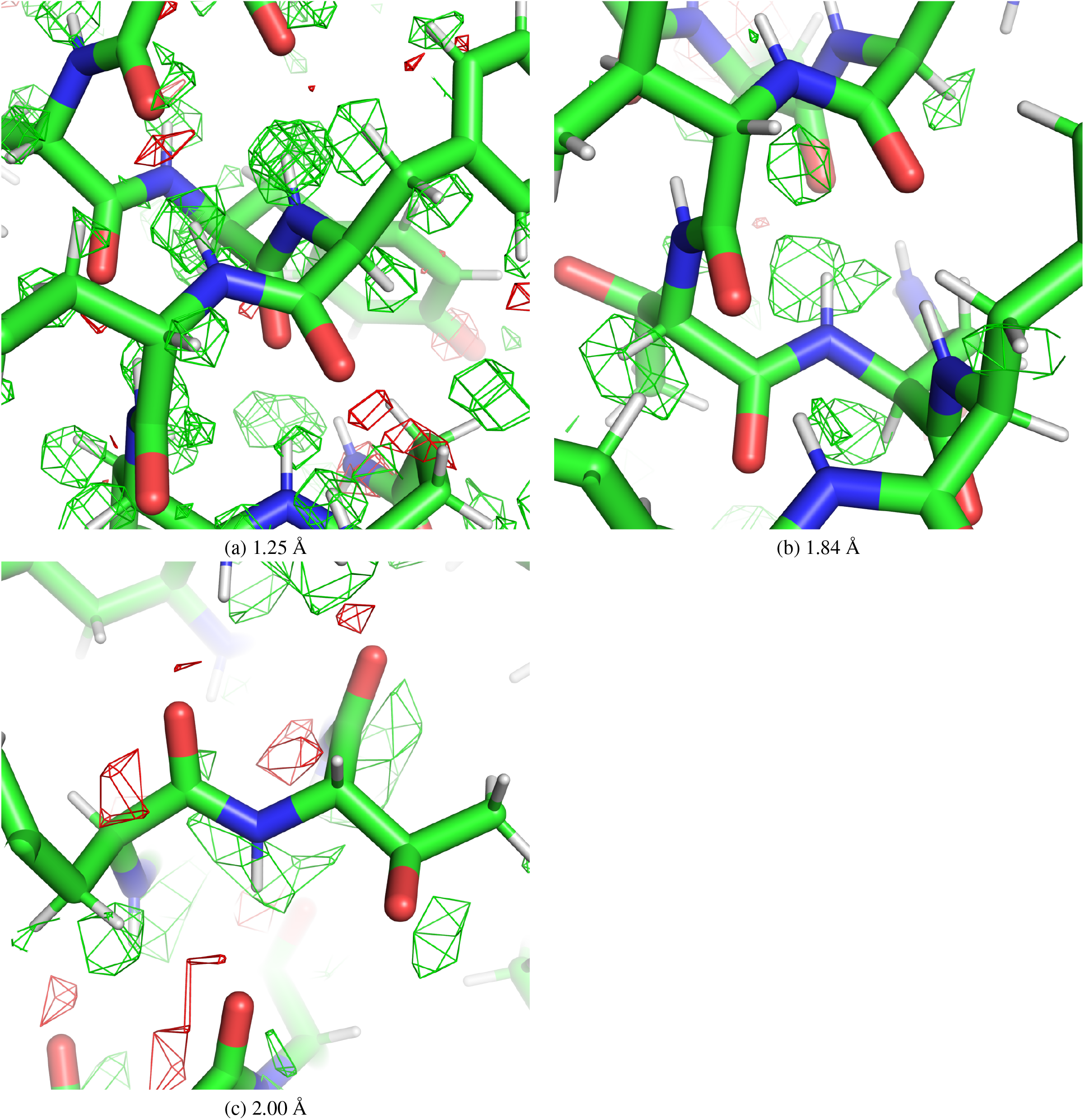
Observation of hydrogen density peaks in *F*_o_ − *F*_c_ maps with different resolutions, using (a) 1.25 Å data (PDB 6z6u/EMD-11103) (b) 1.84 Å data (PDB 6s61/EMD-10101) and (c) 2.00 Å data (PDB 6wx6/EMD-21951). Hydrogen atoms are omitted in the map calculation. Green and red meshes represent positive and negative *F*_o_ − *F*_c_ maps contoured at *±*3*σ* (scaled within the mask), respectively. The images were created using PyMOL.

In addition, *F*_o_ − *F*_c_ maps were made from the 1.2 Å data (PDB 7a4m; EMD-11638) using several different resolution cutoffs. These were analysed in the same way (Fig. 4c), along with *F*_c_ maps calculated from the 7a4m model at the same resolutions (Fig. 4d). Fig. 4c,d show that if the cryo-EM experiment and atomic model refinement are carried out carefully, with due attention to ADPs, then some hydrogen atoms can be seen even at 2.0 Å resolution.

For comparison, we performed the same analysis using X-ray crystallographic data of (apo)ferritins deposited in the PDB. 51 re-refined atomic models available at *PDB-REDO* (Joosten *et al*., 2012) were downloaded, crystallographic *mF*_o_ − *DF*_c_ maps were calculated using *REFMAC*5 and density peaks for hydrogen atoms were analysed as just described. The result (Fig. 4b) confirms that, as expected, hydrogen atoms are more visible in EM than X-ray.

## 6. Conclusions

A new program *Servalcat* for refinement and validation of atomic models using cryo-EM SPA maps has been developed. The program controls the refinement flow and does difference map calculations. A weighted and sharpened *F*_o_ − *F*_c_ map was derived, as a validation tool, obtained from the posterior distribution of *F*_T_ and an approximation of an overall blurring factor calculated from the variance of the signal. We showed that such maps are useful to visualize hydrogen atoms and model errors, as in crystallography. We assumed the blurring factor *k* was position independent. However in reality, blurring of maps is position and direction dependent, e.g. due to varying mobility of different domains and/or uncertainty in the particle alignments. For such regions *k* should be replaced with *k*_local_, where 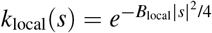 (if isotropic) or 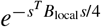 (if anisotropic). If we can estimate *B*_local_ values, then we can use them for visual improvement of maps. We are working on this subject.

We showed that many hydrogen atoms may be observed in the difference maps, even up to a resolution of 2 Å . We would expect that they should also be visible in electron diffraction (MicroED) experiments. However, high accuracy would be needed in the experiment, data analysis, and model refinement in both MicroED and cryo-EM SPA to achieve this experimentally. For example, the electron dose in cryo-EM experiments is often high enough to cause radiation damage (Hattne *et al*., 2018); hydrogen atoms are known to suffer from radiation damage (Leapman & Sun, 1995) and this would hinder their detection. Lower dose experiments might be needed for for more reliable identification of hydrogen, even at the expense of resolution.

Symmetry is widely used in cryo-EM SPA. When symmetry is imposed in the reconstruction, it should be used throughout the downstream analyses, and all software tools should be aware of it and take it into account. The asymmetric unit model should be refined under symmetry constraints, and it should be deposited to the PDB with correct annotation of the symmetry. The PDB and EMDB deposition system will need to validate the symmetry of both model and map. We hope this will become a common practice in future. The same practice should be established for helical reconstructions, in which symmetry is described by the axial symmetry type (*Cn* or *Dn*), twist, and rise (He & Scheres, 2017). *Servalcat* will support helical symmetry in future.

*Servalcat* is freely available under an open source (MPL-2.0) licence at https://github.com/keitaroyam/servalcat. Features described in this paper have been implemented in *REFMAC*5.8.0272 and *Servalcat* 0.2.0 (which requires *GEMMI* 0.4.7). *Servalcat* is also available in the latest nightly builds of the *CCP-EM* suite and will be included in the upcoming v1.6 release.

## Appendix A Variance of a masked map

In this Appendix we demonstrate how a mask inflates sigma-scaled density and show that it is useful to normalize the map using the standard deviation within the mask.

We consider a masked map containing *n* points in total, where *m* points are within the mask and thus the values for *n* − *m* points are zero. If we calculate the mean value of the whole data:

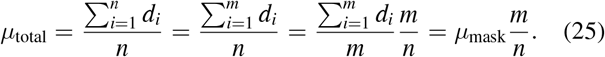

Thus, to calculate the mean within the mask we can calculate the total mean and then use the formula for correction:

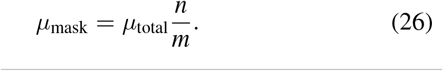

For the variance,

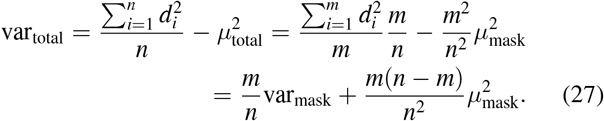

From here we can calculate var_mask_ if we know var_total_ and *µ*_total_. If we denote *f* = *m*/*n* then we can write:

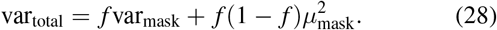

If the mean inside the mask is zero then there is a simple relationship between total variance and variance within the mask. This explains the dependence between the box size and the r.m.s.d. of a cryo-EM SPA map. *Servalcat* normalizes the *F*_*o*_ − *F*_*c*_ map by 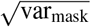 using a user-supplied mask.

If we assume that the map consists of signal and noise, and there is no correlation between them then we can claim that var_mask_ = var_signal_ + var_noise_. Now, in addition, if we assume that we have modelled the map fully with an atomic model (or two maps have almost perfect overlap of signals) then the difference maps should consist almost entirely of noise. Therefore, var_diffmap,mask_ = var_noise_. This variance should be calculated within the mask to make sure that we do not have variance reduction because of systematically low values outside the region occupied by the macromolecule. If we want to increase the reliability of these variances for a region of interest then we may also mask out other regions where there might be signal that is not fully accounted for by the current model. This can be practiced in crystallography as well.

This work was supported by the Medical Research Council, as part of UK Research and Innovation (MC UP A025 1012 to K.Y. and G.N.M.; MR/V000403/1 to C.M.P. and T.B.). The authors are grateful to Marcin Wojdyr for the implementation of *F*_c_ calculation for EM in the *GEMMI* library; Takanori Nakane for critical reading of the manuscript; computational structural biology group members for discussion; Jake Grimmett and Toby Darling from the MRC-LMB Scientific Computing Department for computing support and resources.

## 1. Notation

*F*_T_: Fourier transform of unknown true map (complex values)
*F*_n_: Fourier transform of noise in the observed map (complex values)
*F*_o1_, *F*_o2_: Fourier transforms of the two unweighted and unsharpened half maps from independent reconstructions (complex values)
*F*_o_: Fourier transform of the observed full map: (*F*_o1_ + *F*_o2_)/2
*F*_c_: Fourier transform of calculated map from atomic coordinates (complex values)
*E*: Structure factors normalized in resolution bins: 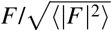
*k*: Resolution dependent scale factor between *F*_o_ and *F*_T_
*D*: Resolution dependent scale factor between *F*_o_ and *F*_c_
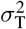: Variance of signal: var(*F*_T_)
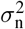: Variance of noise: var(*F*_n_)
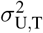: Variance of unexplained signal: var(*DF*_c_ − *kF*_T_)
*f*: Atomic scattering factor
*s*: Column vector of position in reciprocal space
*s*^*T*^: Row vector of position in reciprocal space
*x*: Column vector of position in real space
(*R, t*): Rotation matrix and translation vector that could be an element of a point group Unless otherwise stated, all quantities in Fourier space are dependent on *s*.

## Footnotes

1 There is a similar record, `BIOMT`, which encodes the biological assembly. In SPA, the symmetry of the map usually corresponds to the biological assembly, but this is not always the case. Both `MTRIX` and `BIOMT` records are generally required during the deposition.

2 Technically, fixed position constraints would be more appropriate here. We used restraints instead of constraints for simplicity of implementation. In future, we will implement the use of constraints instead.

