## Supplementary Figures for "Cryo-EM single particle structure refinement and map calculation using *Servalcat*"

Fig. S1: Comparison of weighting and sharpening scheme for Fig. 2a (PDB 7kjr/EMD-22898 at 2.08 Å). (a) Weighted and sharpened map using (17). (b) No FSC-based weights, and sharpening by $B$ value determined by PostProcess in RELION; $(F_\mathrm{o} - DF_\mathrm{c})/e^{-B|s|^2/4}$ with $B = -34.5$ Å$^2$. (c) Using postprocess.mrc; $\sqrt{\mathrm{FSC_{full}}}(F_\mathrm{o} - DF_\mathrm{c})/e^{-B|s|^2/4}$ with the same $B$. The $F_\mathrm{o} - F_\mathrm{c}$ omit maps are contoured at $3\sigma$ (scaled within the mask). The ligand molecule shown as sticks are omitted in the map calculation.

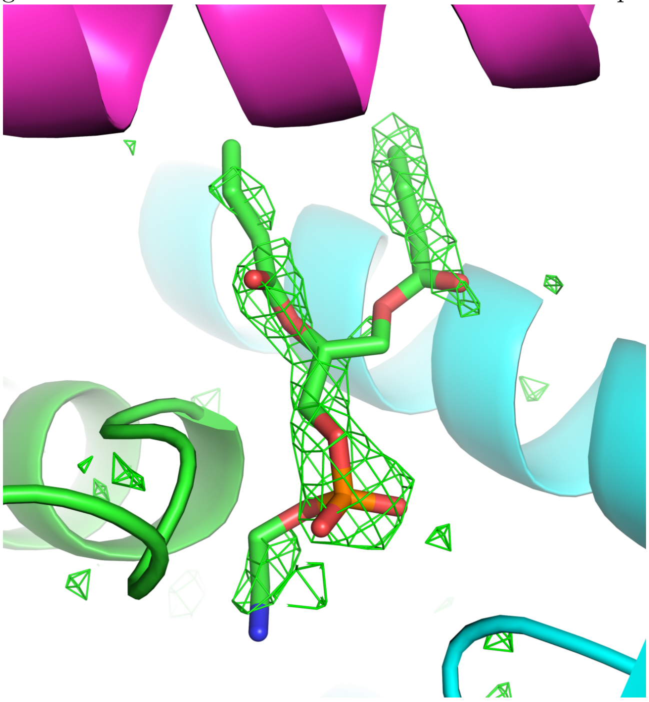

(a) *Servalcat*

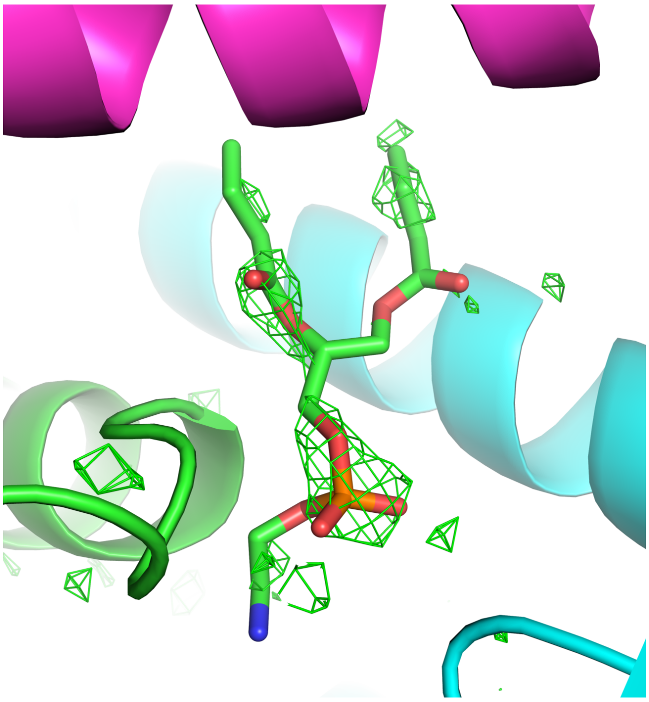

(b) sharpened by $B_\mathrm{PostProcess}$ (no weights)

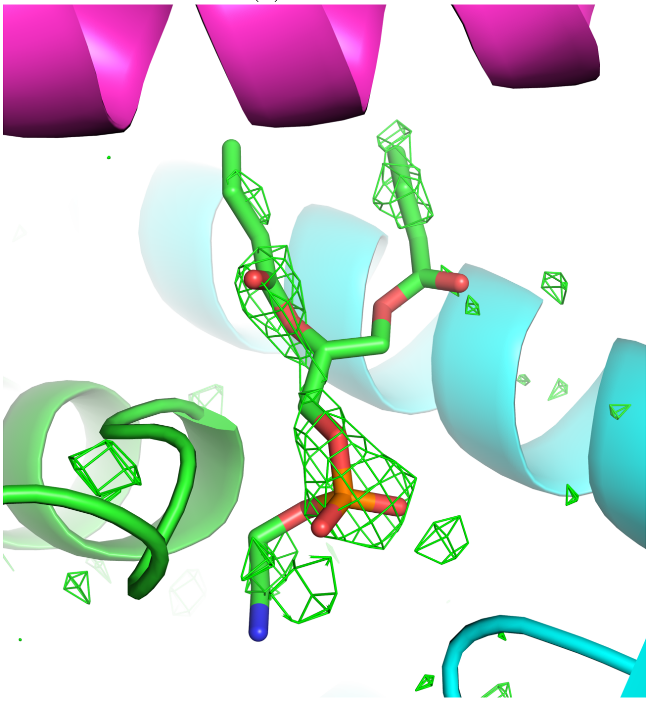

(c) FSC weighted, sharpened by $B_\mathrm{PostProcess}$

Fig. S2: Comparison of weighting and sharpening scheme for Fig. 2b (PDB 5it7/EMD-8123 at 3.6 Å). (a) Weighted and sharpened map using (17). (b) No FSC-based weights, and sharpening by $B$ value determined by PostProcess in RELION; $(F_\mathrm{o} - DF_\mathrm{c})/e^{-B|s|^2/4}$ with $B = -150.1$ Å$^2$. (c) Using postprocess.mrc; $\sqrt{\mathrm{FSC}_\mathrm{full}}(F_\mathrm{o} - DF_\mathrm{c})/e^{-B|s|^2/4}$ with the same $B$. The $F_\mathrm{o} - F_\mathrm{c}$ omit maps are contoured at $3\sigma$ (scaled within the mask). The ligand molecules and ions shown as sticks and spheres, respectively, are omitted in the map calculation.

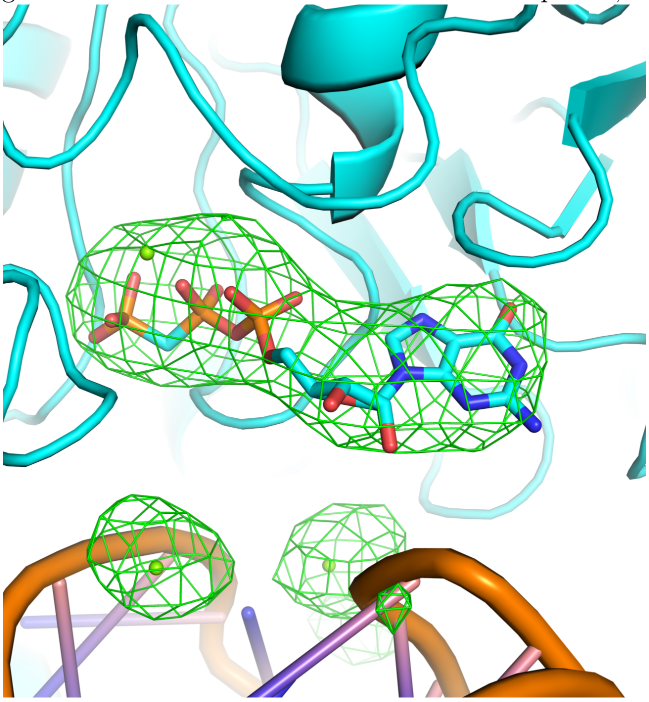

(a) *Servalcat*

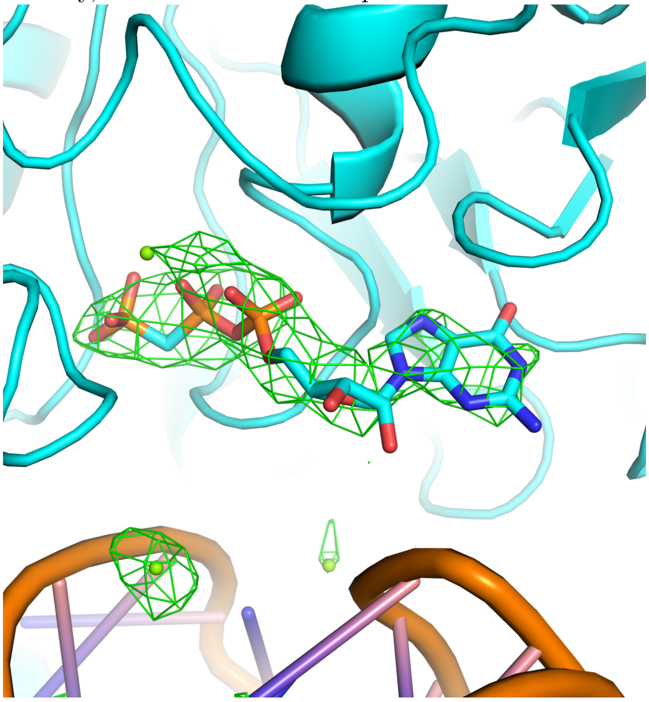

(b) sharpened by $B_\mathrm{PostProcess}$ (no weights)

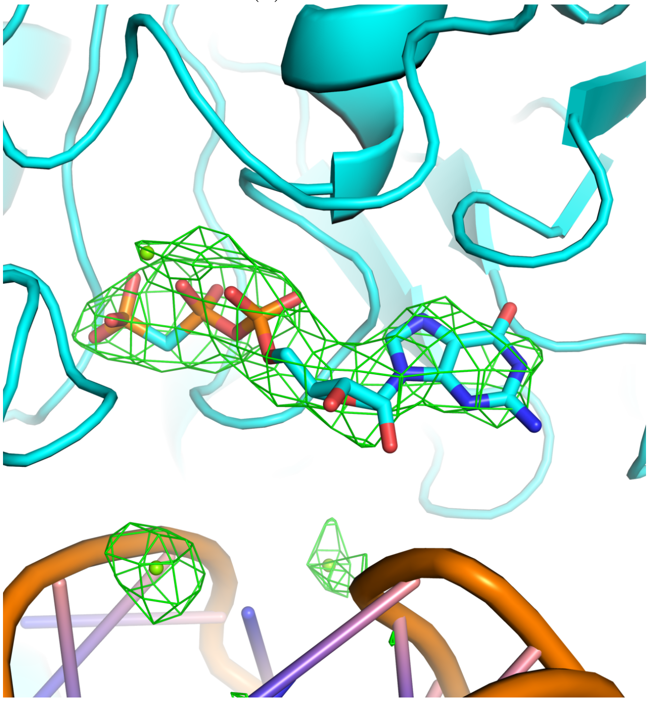

(c) FSC weighted, sharpened by $B_\mathrm{PostProcess}$

Fig. S3: Comparison of weighting and sharpening scheme for Fig. 3 (PDB 6lmt/EMD-0919 at 2.66 Å). (a) Weighted and sharpened maps using (18) and (17). (b) No FSC-based weights, and sharpening by $B$ value determined by PostProcess in RELION; $F_\mathrm{o}/e^{-B|s|^2/4}$ and $(F_\mathrm{o}-DF_\mathrm{c})/e^{-B|s|^2/4}$ with $B=-74.0$ Å$^2$. (c) Using postprocess.mrc; $\sqrt{\mathrm{FSC_{full}}}F_\mathrm{o}/e^{-B|s|^2/4}$ and $\sqrt{\mathrm{FSC_{full}}}(F_\mathrm{o}-DF_\mathrm{c})/e^{-B|s|^2/4}$ with the same $B$. The $F_\mathrm{o}-F_\mathrm{c}$ maps are contoured at $\pm 4\sigma$ (scaled within the mask). The contouring levels of $F_\mathrm{o}$ maps are adjusted to give similar appearance to (a).

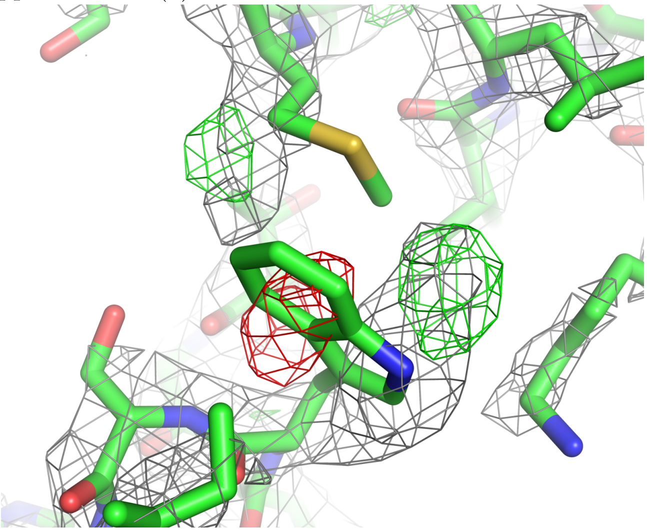

(a) *Servalcat*

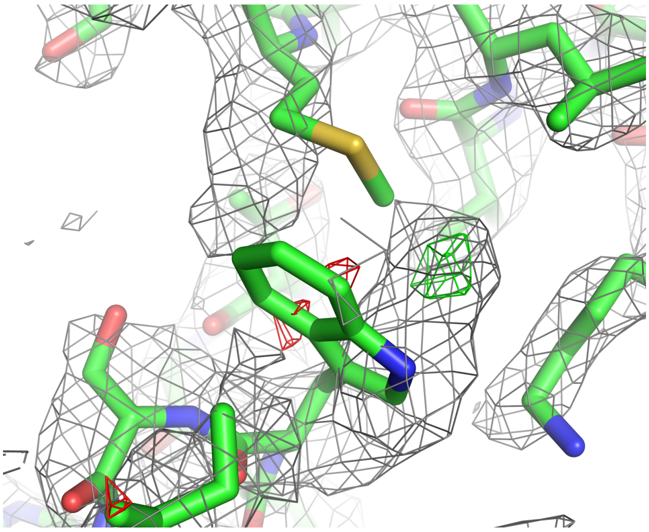

(b) sharpened by $B_\mathrm{PostProcess}$ (no weights)

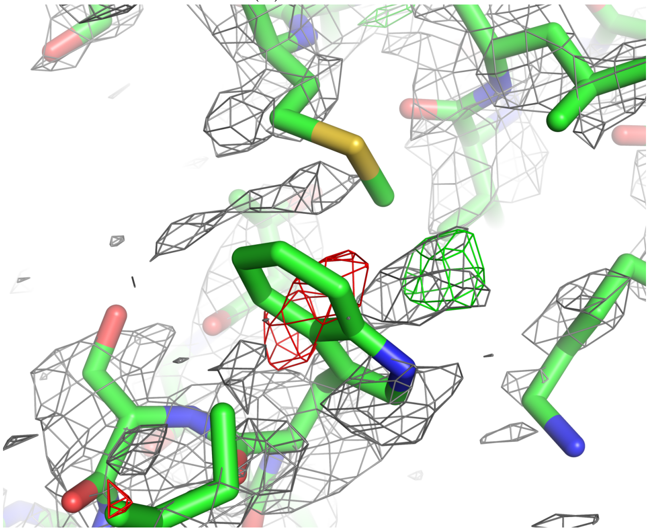

(c) FSC weighted, sharpened by $B_\mathrm{PostProcess}$

Fig. S4: Comparison of weighting and sharpening scheme for Fig. 5a (PDB 6z6u/EMD-11103 at 1.25 Å). (a) Weighted and sharpened map using (17). (b) No FSC-based weights, and sharpening by $B$ value determined by PostProcess in RELION; $(F_\mathrm{o} - DF_\mathrm{c})/e^{-B|s|^2/4}$ with $B = -33.4$ Å$^2$. (c) Using postprocess.mrc; $\sqrt{\mathrm{FSC_{full}}}(F_\mathrm{o} - DF_\mathrm{c})/e^{-B|s|^2/4}$ with the same $B$. The $F_\mathrm{o} - F_\mathrm{c}$ maps are contoured at $\pm 3\sigma$ (scaled within the mask).

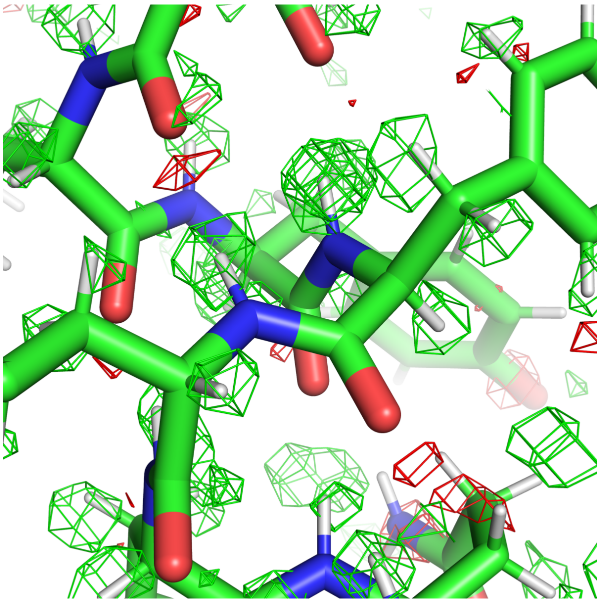

(a) *Servalcat*

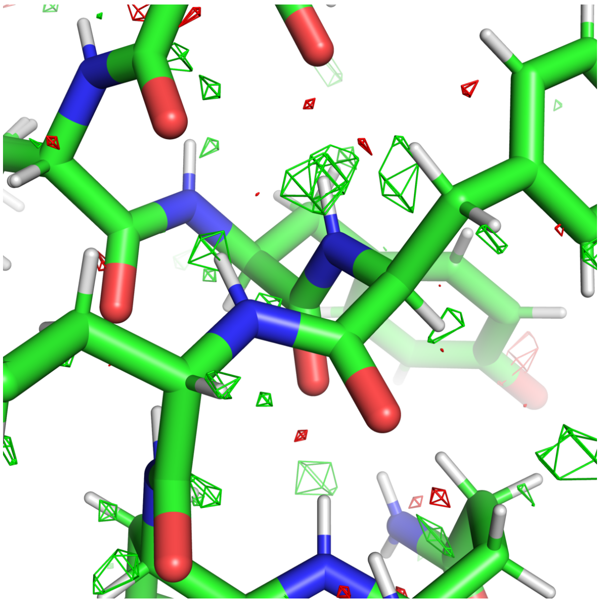

(b) sharpened by $B_\mathrm{PostProcess}$ (no weights)

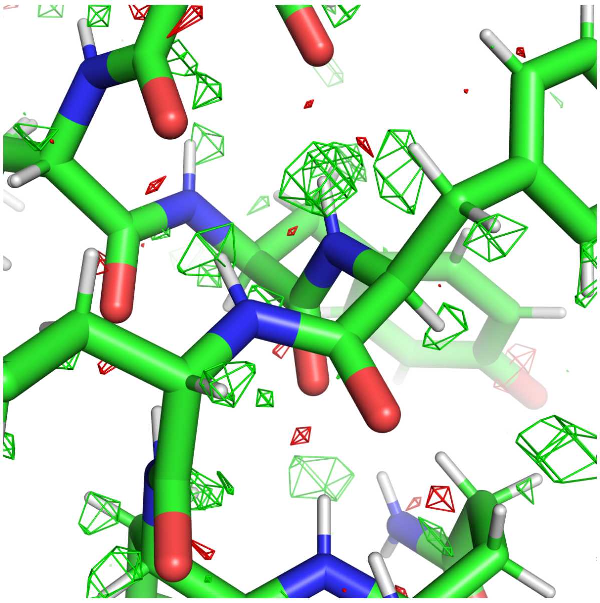

(c) FSC weighted, sharpened by $B_\mathrm{PostProcess}$

Fig. S5: Comparison of weighting and sharpening scheme for Fig. 5b (PDB 6s61/EMD-10101 at 1.84 Å). (a) Weighted and sharpened map using (17). (b) No FSC-based weights, and sharpening by $B$ value determined by PostProcess in RELION; $(F_\mathrm{o} - DF_\mathrm{c})/e^{-B|s|^2/4}$ with $B = -70.9$ Å$^2$. (c) Using postprocess.mrc; $\sqrt{\mathrm{FSC_{full}}}(F_\mathrm{o} - DF_\mathrm{c})/e^{-B|s|^2/4}$ with the same $B$. The $F_\mathrm{o} - F_\mathrm{c}$ maps are contoured at $\pm 3\sigma$ (scaled within the mask).

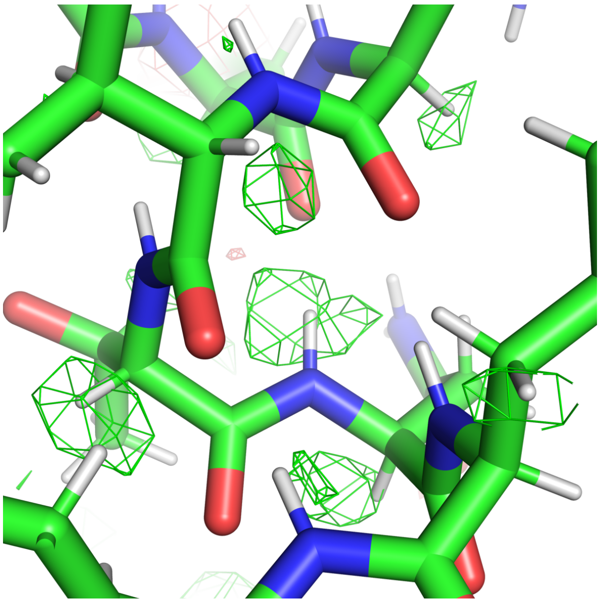

(a) *Servalcat*

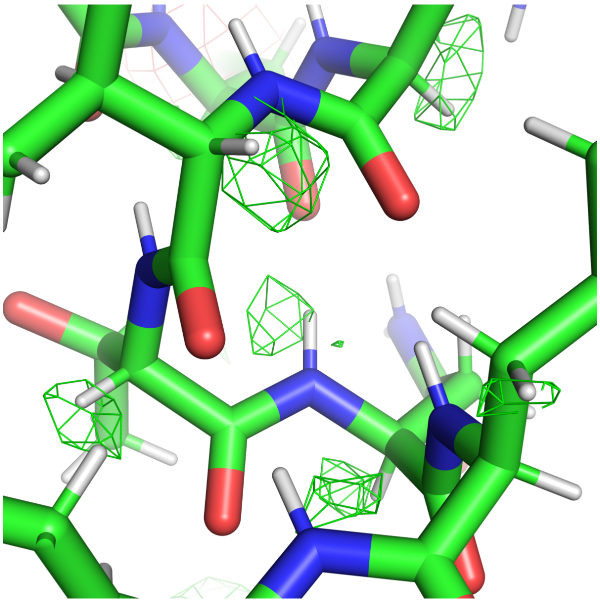

(b) sharpened by $B_\mathrm{PostProcess}$ (no weights)

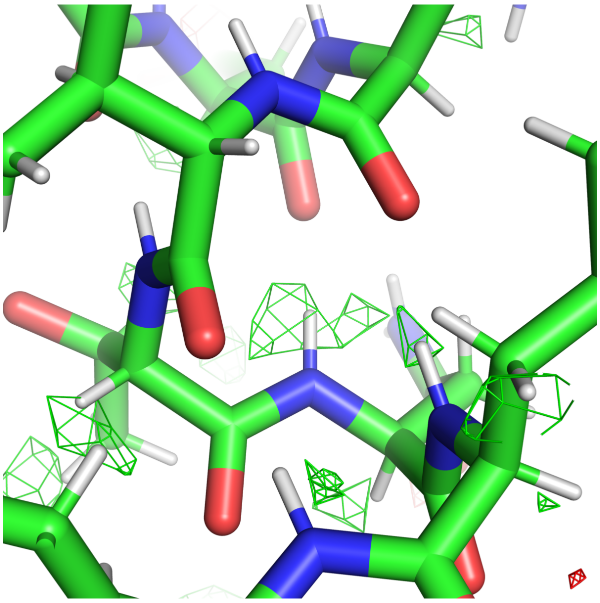

(c) FSC weighted, sharpened by $B_\mathrm{PostProcess}$

Fig. S6: Comparison of weighting and sharpening scheme for Fig. 5c (PDB 6wx6/EMD-21951 at 2.00 Å). (a) Weighted and sharpened map using (17). (b) No FSC-based weights, and sharpening by $B$ value determined by PostProcess in RELION; $(F_\mathrm{o} - DF_\mathrm{c})/e^{-B|s|^2/4}$ with $B = -98.6$ Å$^2$. (c) Using postprocess.mrc; $\sqrt{\mathrm{FSC_{full}}}(F_\mathrm{o} - DF_\mathrm{c})/e^{-B|s|^2/4}$ with the same $B$. The $F_\mathrm{o} - F_\mathrm{c}$ maps are contoured at $\pm 3\sigma$ (scaled within the mask).

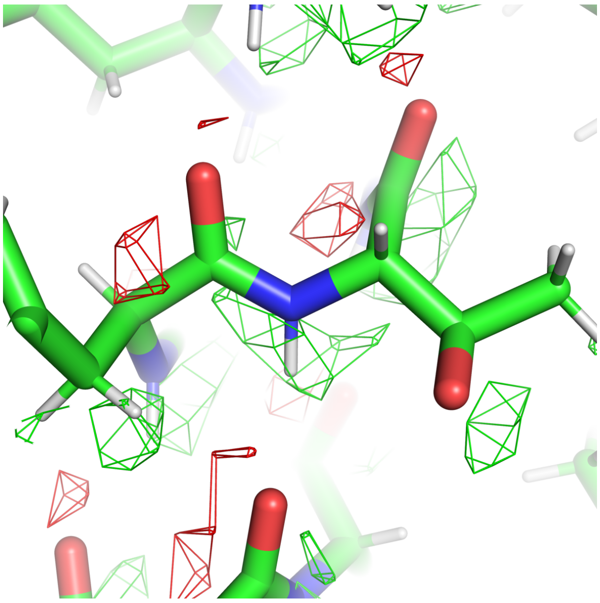

(a) *Servalcat*

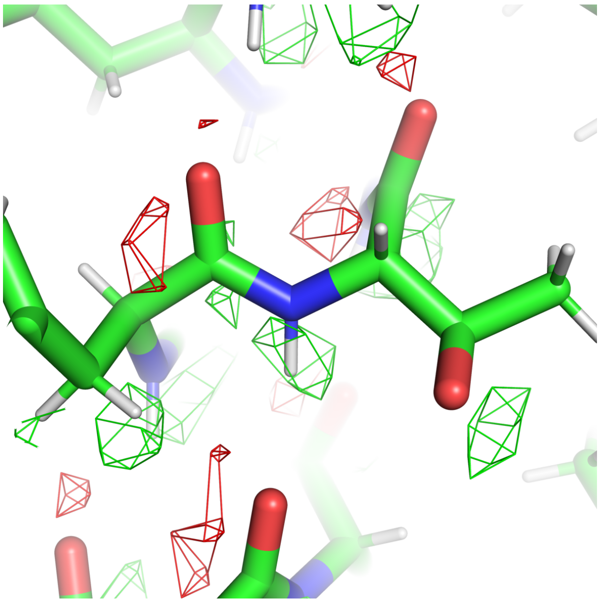

(b) sharpened by $B_\mathrm{PostProcess}$ (no weights)

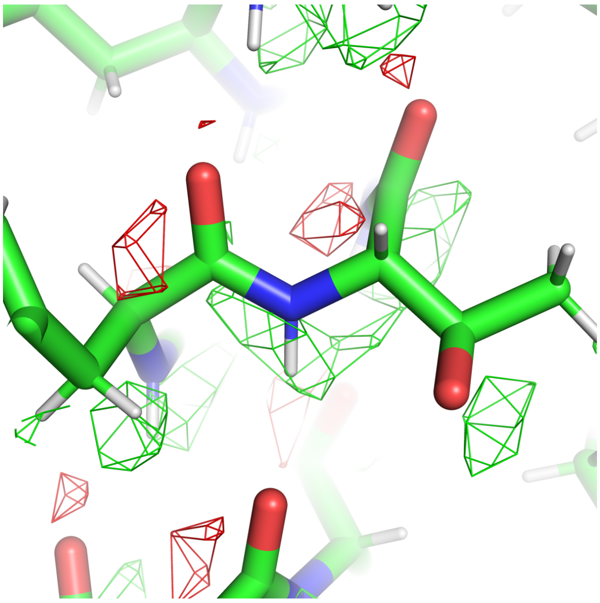

(c) FSC weighted, sharpened by $B_\mathrm{PostProcess}$
